# T-cell activation and HLA-regulated response to smoking in the deep airways of patients with multiple sclerosis

**DOI:** 10.1101/054502

**Authors:** Johan Öckinger, Michael Hagemann-Jensen, Susanna Kullberg, Benita Engvall, Anders Eklund, Johan Grunewald, Fredrik Piehl, Tomas Olsson, Jan Wahlstrom

## 1 Introduction

Multiple sclerosis (MS) is an autoimmune, demyelinating disease of the central nervous system (CNS) [1], with a largely unknown aetiology. International efforts have defined over 100 genetic loci modifying the risk for MS [2]. However, these genes only explain a fraction of the disease risk, suggesting an important contributory role of environmental factors [3]. One of the most established environmental risk factors for MS is cigarette smoking, with an estimated odds ratio (OR) of 1.51 for daily smokers compared to nonsmokers [4]. This risk factor is also shared with other autoimmune diseases, including rheumatoid arthritis (RA), where a risk increase of a similar magnitude is seen [5]. Furthermore, in both MS and RA, there is a significant gene-environment interaction between cigarette smoking and the established HLA-alleles associated with the respective diseases. In MS, the risk is further increased, up to 16-fold, in smokers who carry the risk allele DRB1*1501 in absence of HLA-A*02 [6]. A similar pattern, but with lower overall risk increase, is seen for individuals exposed to second hand smoke [7, 8]. In addition, smokers diagnosed with MS, who continue to smoke after diagnosis display a worsened disease course, with an earlier conversion to a secondary progressive disease course [9], compared to patients who quit smoking after diagnosis. In contrast, the use of moist snuff confers a modest protective effect for the development of MS [10, 11]. Collectively this suggests that exposure, presumably in the lung, to smoke-related compounds rather than nicotine itself, mediates the risk increase, even though the biological mechanisms remain to be established.

Still, a study of the animal MS model experimental autoimmune encephalomyelitis (EAE) in rats demonstrated that transferred autoreactive T-cells migrate to the lungs, and there acquire a migratory phenotype that allow the cells to enter the CNS and initiate an autoimmune disease [12]. This indicates a role for the lung as an immunomodulatory organ, and suggests that similar mechanisms may be involved in the development of human autoimmune disease. We therefore hypothesize that cigarette smoke exposure modifies the function and activation of the immune system, both in the lung and the periphery, and thereby contribute to development and progression of MS. To investigate this we have undertaken the first systematic investigation of the lungs and the pulmonary immune response in smokers and non-smokers diagnosed with MS.

## 2 Material and methods

### 2.1 Study subjects

Healthy volunteers (n=55) were recruited by public advertising, and individuals diagnosed with MS (n=26] were recruited via the Neurology Clinic, Karolinska University Hospital, Stockholm, Sweden. Data from additional healthy controls (n=70) was retrieved from an in-house data registry. Written informed consent was obtained from all subjects, and the Regional Ethical Review Board in Stockholm approved the studies.

None of the subjects had clinically relevant airway infections or allergy symptoms at the time of bronchoscopy, and subjects diagnosed with asthma, COPD, other lung diseases, or other inflammatory conditions were not included in the study. Questionnaires regarding general and pulmonary health, current medications, and historic and current smoking habits were filled out by all subjects. Based on the smoking history we defined *Smokers* as current daily smokers with ≥5 pack years (PY: [cigarettes smoked per day]/20 x [years smoking]) or currently smoking ≥5 cigarettes/day. Previous smokers (≥ 5 PY,≥5 cigarettes/day], who quit ≤24 months before bronchoscopy were also classified as *Smokers* (3 individuals, mean time since cessation: 12 months). *Nonsmokers* were defined as never smokers or previous smokers who quit >5 years before bronchoscopy (1 MS-NS, smoke cessation 20 years before bronchoscopy). Additional healthy controls (Supplementary Table 2) were self-defined as smokers or non-smokers. Information on MS disease type, clinical status at time of bronchoscopy, magnetic resonance imaging (MRI) findings and treatment history (Table 2) was retrieved from clinical records and the Swedish MS registry. All patients with MS fulfilled the McDonald criteria [13]. The patient cohort represented a cross-sectional sample of patients with established disease, reflecting different types of disease course and treatments, however, with a lack of very recently diagnosed treatment naive patients.

The median age of subjects was 28 (19-62) years, and 32 (39.5%) of the subjects were smokers at the time of sample collection with a median smoke history of 6.5 (1.8-27.0) pack years (PY). The MS-patients included in the study have slightly higher median age, and consequently higher cumulative smoking load (PY) but similar current smoking habits (cigarettes/day) compared to the healthy smoking volunteers.

### 2.2 Spirometry and bronchoscopy with bronchoalveolar lavage (BAL)

All included subjects underwent dynamic spirometry (Jaeger MasterScope, Intramedic; Medikro PRO, Aiolos). Bronchoscopy with bronchoalveolar lavage (BAL) was performed as previously described [14]. In short, bronchoscopies were carried out in the supine position with a flexible fiberoptic bronchoscope (Olympus Optical Co. Ltd, Tokyo, Japan] inserted nasally. The bronchoscope was wedged in a subsegmental bronchus in the middle lobe and 5 aliquots of 50 ml of sterile, phosphate-buffered saline solution at +37°C were instilled. After each instillation the fluid was gently aspirated with a negative pressure of 6-10kPa. The aliquots of bronchoalveolar lavage fluid (BALF) were pooled, collected in a siliconized plastic bottle and kept on ice until further analysis.

### 2.2.1 Differential count of bronchoalveolar lavage (BAL)

Smears for differential cell counts were prepared by cytocentrifugation (Cytospin 2; Shanon Ltd, Runcorn, UK) at 22 g for 3 minutes and stained with May-Grünwald Giemsa. A minimum of 500 cells was counted, and mast cells in 10 visual fields (16 x magnification) were determined after staining with toluidine/haematoxylin.

### 2.3 Isolation of cells from peripheral blood

Peripheral blood mononuclear cells (PBMCs) were isolated using Ficoll-Paque (GE Healthcare, Little Chalfont, UK) separation from heparinized peripheral blood and resuspended in RPMI 1640 supplemented with HEPES, 10% Human Serum, 1% Penicillin-Streptavidin, 1% L-Glutamine (Sigma Aldrich, St. Louis, MO). Both BALF and peripheral blood were obtained from all subjects in the study cohort.

### 2.4 Flow cytometry

Surface staining was performed on *ex vivo* BALF cells and PBMC's washed twice in PBS + 1% human Ab serum (Sigma]. The following antibodies were used: CD4-PerCP-Cy5.5 (SK3), CD3-APC-H7 (SK7), and CD8-V500 (SK1) (all from BD Bioscience Franklin Lakes, NJ). Intracellular staining was performed after fixation and permeabilization of surface stained cells by Anti-Human Foxp3 staining kit (eBiosciences San Diego, CA), using Ki-67-AlexaFlour 488 (B56), CD154-APC (TRAP-1), mouse IgG1-AlexaFlour 488 (MOPC-21), mouse IgG1-APC (MOPC-21) (BD Bioscience). The flow cytometry data acquisition was performed on a BD FACS Canto-II flow cytometer (BD Bioscience) and the analysis was carried out using TreeStar FlowJo 10. All samples were analysed after exclusion of duplets and according to respective isotype-controls.

### 2.5 HLA genotyping

Genomic DNA was extracted from whole blood samples and HLA-DRB1 alleles (two-digit resolution) in healthy subjects were determined using polymerase chain reaction (PCR) amplification with sequence-specific primers (DR low resolution kit, Olerup SSP, Saltsjöbaden, Sweden). HLA-DRB1 alleles in 16 MS patients were imputed from Immunochip project data as previously described [15], with an estimated concordance rate of 99.0% to sequence data, for HLA-DRB1 at two-digit resolution. Nine MS patients were genotyped using TaqMan (Applied Biosystems, Waltham, MA) allelic discrimination of the tag-SNPs rs9271366, with a reported sensitivity and specificity >97% for presence of HLA-DRB1*15 [8]. HLA-DRB1 genotype could not be obtained from one of the MS-patients.

### 2.6 Statistical analysis

Statistics were calculated using GraphPad Prism 5 and 6, or R 3.1.2. Statistical methods for each analysis are indicated in the main text or figure legends. p<0.05 was considered statistically significant.

## 3 Results

### 3.1 Minor airway obstruction observed in MS-patients

We recruited 81 patients and volunteers for this study, who all underwent clinical assessment, spirometry and bronchoscopy with bronchoalveolar lavage. Interestingly, the MS patients showed an overall reduced FEV1/FVC ratio compared to the healthy subjects independently of smoking status (Table 1), gender and smoking habits (cigarettes/day) or cumulative smoking (PY) (data not shown), but modified by age (Supplementary Table 1). This might indicate a subclinical process affecting the airways in subgroups of MS patients. The clinical characteristics of patients with MS was also analysed based on current smoking status (Table 2). No differences in clinical parameters between smokers and non-smokers could however be detected in this study.

**Table 2:** Clinical characteristics and current treatment, in MS-patients. Disease duration: median time (years) and range since disease onset, OCB: occurrence of oligoclonal bands in cerebrospinal fluid, MR lesions: number of inflammatory lesions in CNS detected by magnetic resonance imaging, EDSS: median and range of Extended Disability Status Scale score, PP: primary progressive, RR: relapsing remitting, SP: secondary progressive, relapse/remission indicated for RR patients only. DMT: Disease modifying treatments in MS; Un: untreated. IFN: Interferon-β (Avonex (n=9) or Betaferon (n=2)). Na: Natalizumab, O: Other (including Glatiramer (n=1), Rituximab (n=1), Fingolimod (n=2)). Statistics calculated using Mann Whitney, Chi-2 and Fischer's exact tests, as applicable.

|  | Number | Duration<br>(years) | OCB<br>-/+ | MR lesions<br>(1-9/10-20/>20) | EDSS | Disease course<br>(PP/RR/SP) | Relapse/<br>remission | DMT<br>(Un/IFN/Na/O) |
| --- | --- | --- | --- | --- | --- | --- | --- | --- |
| **MS total** | 26 | 5 (0-20) | 3/23 | 6/11/9 | 2.3 (0-5) | 2/21/3 | 5/16 | 6/11/5/4 |
| **Smokers** | 11 | 4 (1-20) | 1/10 | 3/5/3 | 2.5 (0-5) | 0/11/0 | 2/9 | 3/4/3/1 |
| **Non-smokers** | 15 | 5 (0-9) | 2/13 | 3/6/6 | 2.0 (1-3.5) | 2/10/3 | 3/7 | 3/7/2/3 |

**Table 1:** Characterization of patients and volunteers included in the study. Numbers represent median and (range) if not otherwise stated. Statistics calculated using Kruskal-Wallis test with Dunn's post test: ^a^ p<0.001 compared to healthy non-smokers Statistics calculated using Mann Whitey (2 groups): ^b^ p<0.01, compared to healthy smokers. Statistics calculated using linear regression analysis for independent effects of smoke status and diagnosis: ^#^ p<0.05 compared to healthy volunteers (smoke status not significant) ^1^ Pack years (PY: [cigarettes smoked per day]/20 x [years smoking]). ^2^ current smoking habits at bronchoscopy. ^3^ FVC: forced vital capacity, % of predicted. ^4^ FEV1: Forced expiratory volume in 1 second, % of predicted. n.a: not applicable.

| Characteristics | Healthy volunteers |  | Multiple Sclerosis |  |
| --- | --- | --- | --- | --- |
|  | Non-Smokers | Smokers | Non-Smokers | Smokers |
| **Total number** | 34 | 21 | 15 | 11 |
| **Gender (M/F)** | 16/18 | 6/15 | 5/10 | 2/9 |
| **Age (years)** | 23.5 (20-57) | 25 (19-50) | 42 (30-62)<sup>a</sup> | 38 (24-55) |
| **Pack Years<sup>1</sup>** | n.a. | 5 (1.8-24.5) | n.a. | 13 (6-27)<sup>b</sup> |
| **Cigarettes/day<sup>2</sup>** | n.a. | 10 (5-30) | n.a. | 10 (1-20) |
| **FVC, liter<sup>3</sup>** | 105.5 (86-128) | 107 (84-127) | 102 (80-137) | 108 (91-137) |
| **FEV1, liter<sup>4</sup>** | 103.5 (82-122) | 103 (80-121) | 101 (78-123) | 106 (80-124) |
| **FEV1/FVC %** | 84 (72-93) | 80 (75-92) | 83 (55-86)<sup>#</sup> | 81 (70-89)<sup>#</sup> |

### 3.2 Cigarette smoking induces increase of alveolar macrophages

As expected, smokers from both the MS and control groups showed an altered composition of immune cells in the bronchoalveolar lavage fluid (BALF) (Table 3). Overall, the results corresponded to previous analyses of cellular content in the lungs of smokers without signs of smoke-related pathology [16]. The two groups of smokers, healthy volunteers and patients with MS, both had a significant increase in the total number of cells, primarily due to an increase in number of alveolar macrophages. Regression analysis on diagnosis, smoking status and age further revealed that smoking is associated with a reduced recovery rate of BALF. Smoking is also associated with an increase in total cell number, concentration of total cells, concentration and relative frequency of macrophages, but a reduced frequency of lymphocytes in BALF, independent of diagnosis (Table 3 and Supplementary Table 1). In addition, higher age was associated with an increase in total cell yield, concentration of total cells and lymphocytes, as well as small but significant changes in the relative distribution of macrophages and lymphocytes in BALF (Supplementary Table 1).

**Table 3.** BAL characteristics and differential cell count in BALF. Numbers indicate median and range if not otherwise stated. Statistics calculated using Kruskal-Wallis test with Dunn's post test ^a^ p<0.05 compared to healthy non-smokers. ^b^ p<0.001 compared to healthy non-smokers. ^c^ p<0.05 compared to MS non-smokers. Statistics calculated using linear regression analysis for independent effects of smoke status and diagnosis, ^#^ p<0.05, ^##^ p<0.01, ^##^ p<0.001^###^, all compared to non-smokers (diagnosis not significant) ^1^ calculated in 16x magnification

| Characteristics | Healthy volunteers |  | Multiple Sclerosis |  |
| --- | --- | --- | --- | --- |
|  | Non-Smokers | Smokers | Non-Smokers | Smokers |
| Performed BAL. no | 34 | 21 | 15 | 11 |
| Recovery (%) | 74 (41-84) | 70 (25-77) <sup>#</sup> | 64 (27-78) <sup>a, #</sup> | 56 (52-76) |
| Total cell yield (x 10<sup>6</sup>) | 15.3 (7.7-29.0) | 39.8 (15.9-154.0) <sup>b, ##</sup> | 14.4 (7.4-28.9) | 34.0 (8.2-166.0) <sup>c, ##</sup> |
| Cell concentration (x 10<sup>6</sup> /L) | 76.4 (14.3-279.5) | 257.8 (90.3-987.2) <sup>c, ##</sup> | 119.5 (48.8-173.7) | 314.8 (66.4-902.0) <sup>##</sup> |
| ***Differential count of BALF*** |  |  |  |  |
| Macrophages % | 91.7 (61.9-96.6) | 96.2 (61.4-99.0) <sup>b, ##</sup> | 93.0 (72.0-96.8) | 95.4 (80.6-97.4) <sup>##</sup> |
| Macrophages x 10<sup>6</sup> /L | 70.9 (12.7-261.6) | 255.2 (83.1-949.7) <sup>b, ###</sup> | 101.0 (46.6-150.3) | 304.1 (61.9-851.5) <sup>###</sup> |
| Lymphocytes % | 6.0 (2.4-31.6) | 2.2 (0.4-7.8) <sup>b, ##</sup> | 6.0 (2.4-27.2) | 3.0 (1.2-18.0) <sup>##</sup> |
| Lymphocytes x 10<sup>6</sup> /L | 5.5 (1.5-50.3) | 6.5 (1.0-29.6) | 5.4 (1.9-44.0) | 10.5 (1.6-30.1) |
| Neutrophils % | 1.2 (0.2-6.4) | 0.6 (0.2-2.8) | 1.0 (0.2-10.0) | 0.8 (0-3.4) |
| Neutrophils x 10<sup>6</sup> /L | 1.0 (0.1-11.2) | 1.5 (0.4-9.2) | 1.3 (0.1-9.6) | 1.8 (0-30.7) |
| Eosinophils % | 0.0 (0.0-2.3) | 0.0 (0.0-1.4) | 0.0 (0.0-2.0) | 0.0 (0.0-1.0) |
| Eosinophils x 10<sup>6</sup> /L | 0.0 (0.0-4.7) | 0.0 (0.0-3.9) | 0.0 (0.0-1.9) | 0.0 (0.0-4.3) |
| Basophils % | 0.0 (0.0-0.4) | 0.0 (0.0-0.2) | 0.0 (0.0-0.2) | 0.0 (0.0-0.0) |
| Basophils x 10<sup>6</sup> /L | 0.0 (0.0-0.4) | 0.0 (0.0-0.8) | 0.0 (0.0-0.3) | 0.0 (0.0-0.0) |
| Mast cells/10 fields of vision <sup>1</sup> | 1 (0-10) | 1 (0-6) | 1 (0-12) | 0 (0-5) |

### 3.3 Distinct distribution of T-cells in the lungs of smokers and MS-patients

Characterization of the T-cells in BALF was done using flow cytometry. This analysis revealed that smoking was associated with a significant reduction of CD4+ cells in MS smokers, compared to MS-non-smokers, and a significant shift in the CD4/CD8 ratio between smokers and non-smokers among the healthy volunteers (Figure 1).

**Figure 1:**
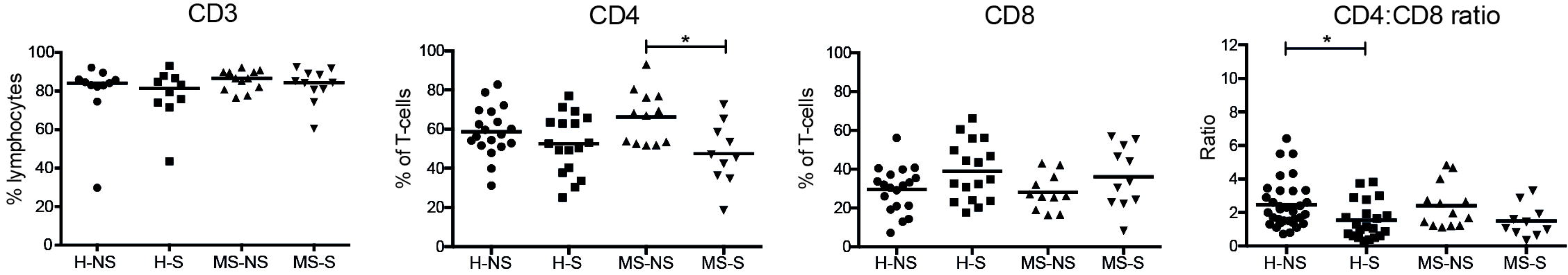
Relative frequency of CD3+, CD4+ and CD8+ T-cells, and CD4+/CD8+ ratio in BALF, analysed by flow cytometry. Y-axes labels indicate which cell subsets were gated on. Horizontal lines indicate median. Statistics calculated using Kruskal-Wallis test with Dunn's post-test, ^*^: p <0.05. H: healthy volunteers, MS: patients diagnosed with MS. NS: non-smokers, S: smokers

The proportion of proliferating T-cells in the BALF was estimated using the marker Ki-67. Both smoking MS-patients and healthy smokers displayed a higher proportion of proliferating (Ki-67+) CD4+ and CD8+ T-cells, compared with healthy non-smokers (Figure 2A). Non-smokers with MS also showed an increase in Ki-67+ cells, compared to healthy non-smokers (Mann Whitney test, CD4+: p=0.017, CD8+: p=0.046), although not significant in the ANOVA analysis. All groups showed however a significant increase in the number of proliferating CD4+ (Figure 2B) and CD8+ (not shown) cells in the BALF, compared to blood.

**Figure 2:**
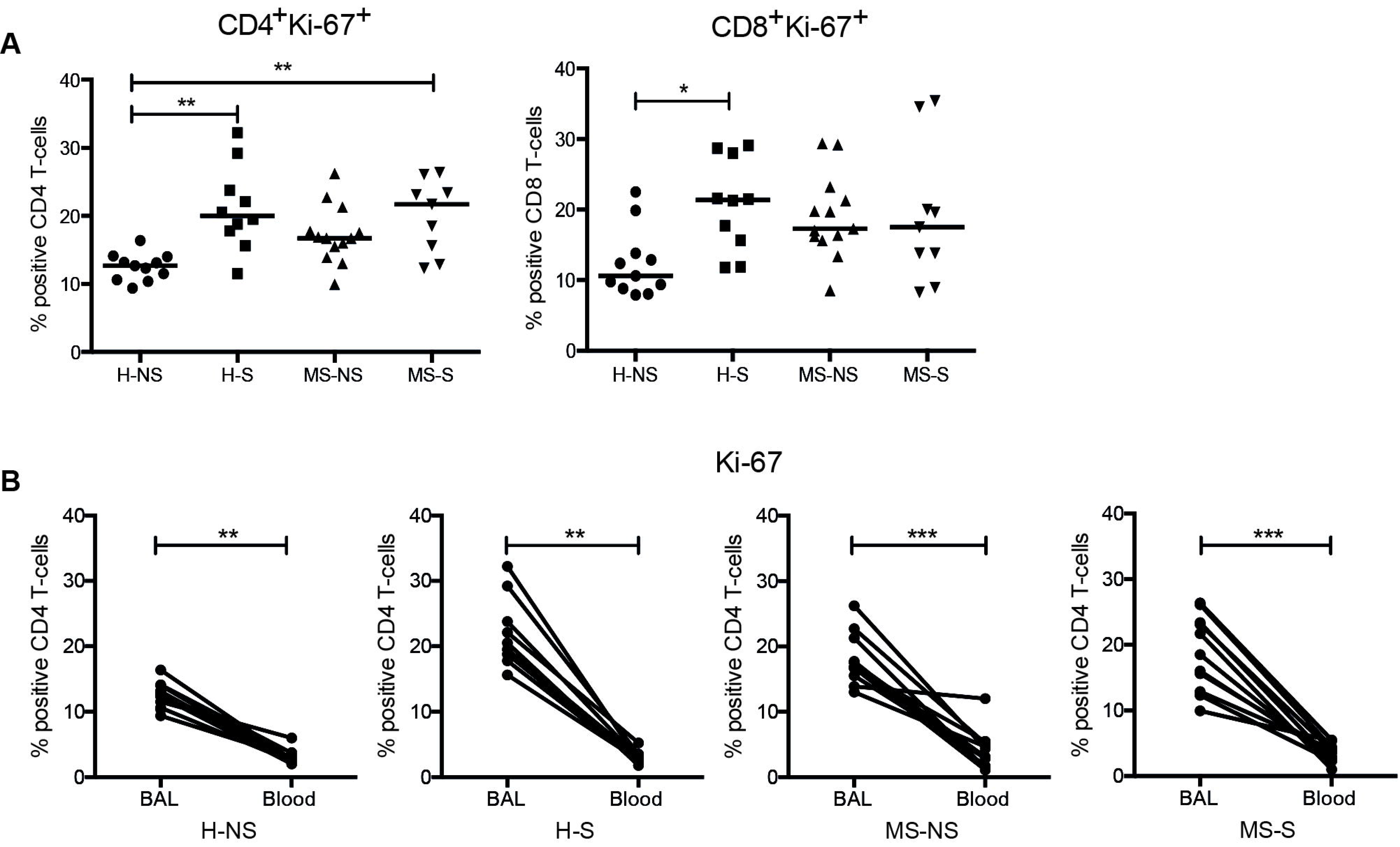
Increased proliferation of T-cells in BALF T-cells. A) Relative frequency of CD4+ KI-67+ and CD8+ KI-67+ T-cells in BALF analyzed by flow cytometry. Statistics calculated using Kruskal-Wallis test with Dunn's post-test, ^*^: p <0.05, ^**^: p <0.01, ^***^: p <0.001. B) Pairwise comparison of frequencies of CD4+ Ki-67+ T-cells in BALF and blood from the same individuals in each respective group. Horizontal lines indicates median, statistics calculated using Wilcoxon matched-pair signed rank test, ^**^: p <0.01, ^***^: p <0.001. H: healthy volunteers, MS: patients diagnosed with MS. NS: non-smokers, S: smokers

CD40L (CD154) is a vital component for optimal CD4+ and CD8+ T-cell responses and licensing of antigen presenting cells (APCs). Dysregulation of CD40L expression and function has been linked to autoimmunity and defective immunity, reviewed in [17]. Here, we analysed the intracellular, preformed CD40L (pCD40L) content *ex vivo* in BALF CD4+ T-cells and identified an increase in pCD40L median fluorescence intensity (MFI) in cells from non-smokers diagnosed with MS, compared to healthy non-smokers (Figure 3A), as well as an increase in frequency of pCD40L+ CD4+ T-cells (Figure 3B). The increase of pCD40L MFI in MS-patients was not significantly regulated by disease modulatory treatment (DMT) status (Supplementary Figure 2). In smokers, we could further see a negative correlation between frequency of pCD40L+ CD4+ T-cells and current smoking habits, with no significant difference between MS and healthy subjects (Figure 3C).

**Figure 3:**
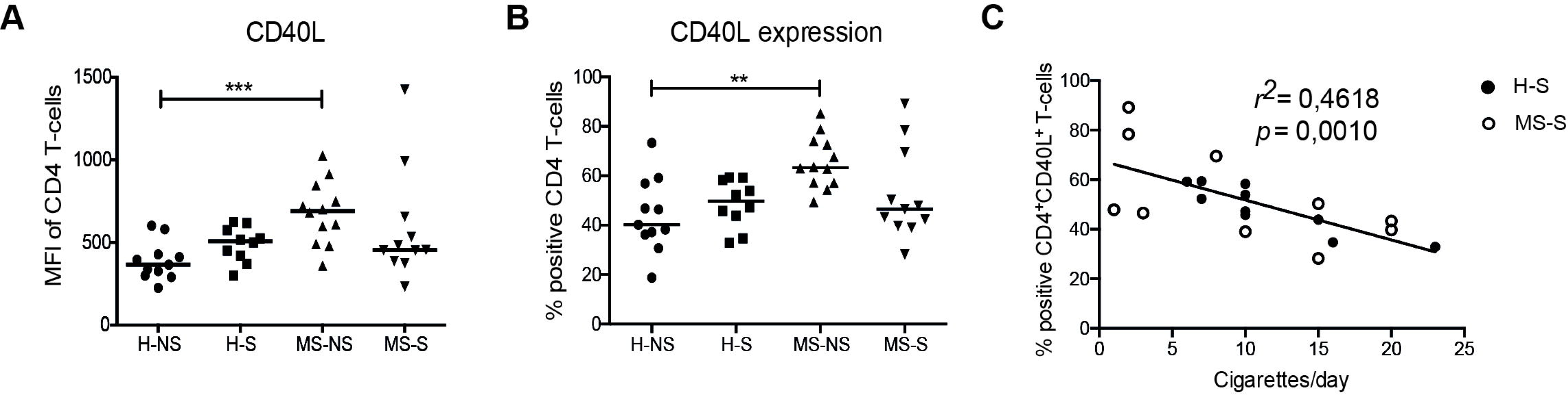
Distinct distribution of intracellular pCD40L in healthy subjects and MS-patients. A) Median fluorescence intensity of intracellular pCD40L in CD4+ T-cells in BALF, and B) frequency of CD40L+, CD4+ T-cells in BALF, measured by flow cytometry Horizontal lines indicate median, statistics calculated using Kruskal-Wallis test with Dunn's post-test, ^**^: p <0.01. ^***^ : p <0.001. C) Spearman rank correlation between CD40L+ CD4+ frequency and current smoking habits (cigarettes/day) in healthy and MS smokers. H: healthy volunteers, MS: patients diagnosed with MS. NS: non-smokers, S: smokers

As age was associated with a significant increase in the concentration of lymphocytes in the BALF, we further analysed the independent effects of diagnosis, smoking status and age on the analysed T-cell subsets and phenotypes, using regression analysis. This analysis confirmed that smoking is associated with a shift in relative frequency of CD4+ and CD8+ T-cells, as well as increased proliferation (Ki-61+), independent of diagnosis, while an MS-diagnosis was significantly associated with increase of pCD40L, independent of smoking status. Age was not significantly associated with any of the analysed phenotypes or subsets of T-cells (Supplementary Table 1).

### 3.4 Cellular distribution in the lung is not associated with treatment

The majority of the MS patients included in this study received either natalizumab or IFN-β as DMT, and also included a group without on going DMT. Cellular distribution and T-cell subsets in BALF were stratified based on DMT status, however not showing any significant differences in the analysed parameters (Table 2Table 2).

### 3.5 HLA-DRB1*15 is associated with attenuated response to cigarette smoke

To investigate if the major genetic risk factor in MS, HLA-DRB1*15, influences the numbers or distribution of immune cells in the lung of smokers and non-smokers, we stratified all subjects by smoking status and HLA-DRB1 allele. As the increased concentration of alveolar macrophages (AM) in the BALF was seen in smokers, regardless of diagnosis (Table 3), both MS-patients and healthy controls were combined in the groups of smokers and non-smokers, respectively. This analysis revealed that smokers carrying HLA-DRB1*15 displayed a reduced concentration of AM compared to non-carriers (Figure 4A), regardless of smoking habits, cumulative smoking or age (not shown), and a similar trend was also seen in MS-patients alone (Figure 4B). In contrast, no association between concentration of AM and HLA-DRB1*15 was seen in the nonsmokers. To further investigate the role of HLA-DRB1 alleles in the response to smoke exposure we also included an additional cohort of healthy volunteers that previously had undergone bronchoscopy with BAL using an identical procedure [18–20] (Supplementary Table 2). Analysis of this extended cohort of healthy individuals confirmed that HLA-DRB1*15 is associated with an attenuated increase of AM concentration in the BALF from smokers, while no difference was seen in non-smokers (Figure 4C). In contrast, we identified that smokers positive for HLA-DRB1*01 or DRB1*03 have an increased concentration of AM in BALF, compared to smokers negative for the respective allele. Other analysed alleles were not significantly associated with concentration of alveolar macrophages in BALF, and none of the HLA-DRB1-alleles were significantly associated with concentration of lymphocytes or other cell types in BALF (data not shown)

**Figure 4:**
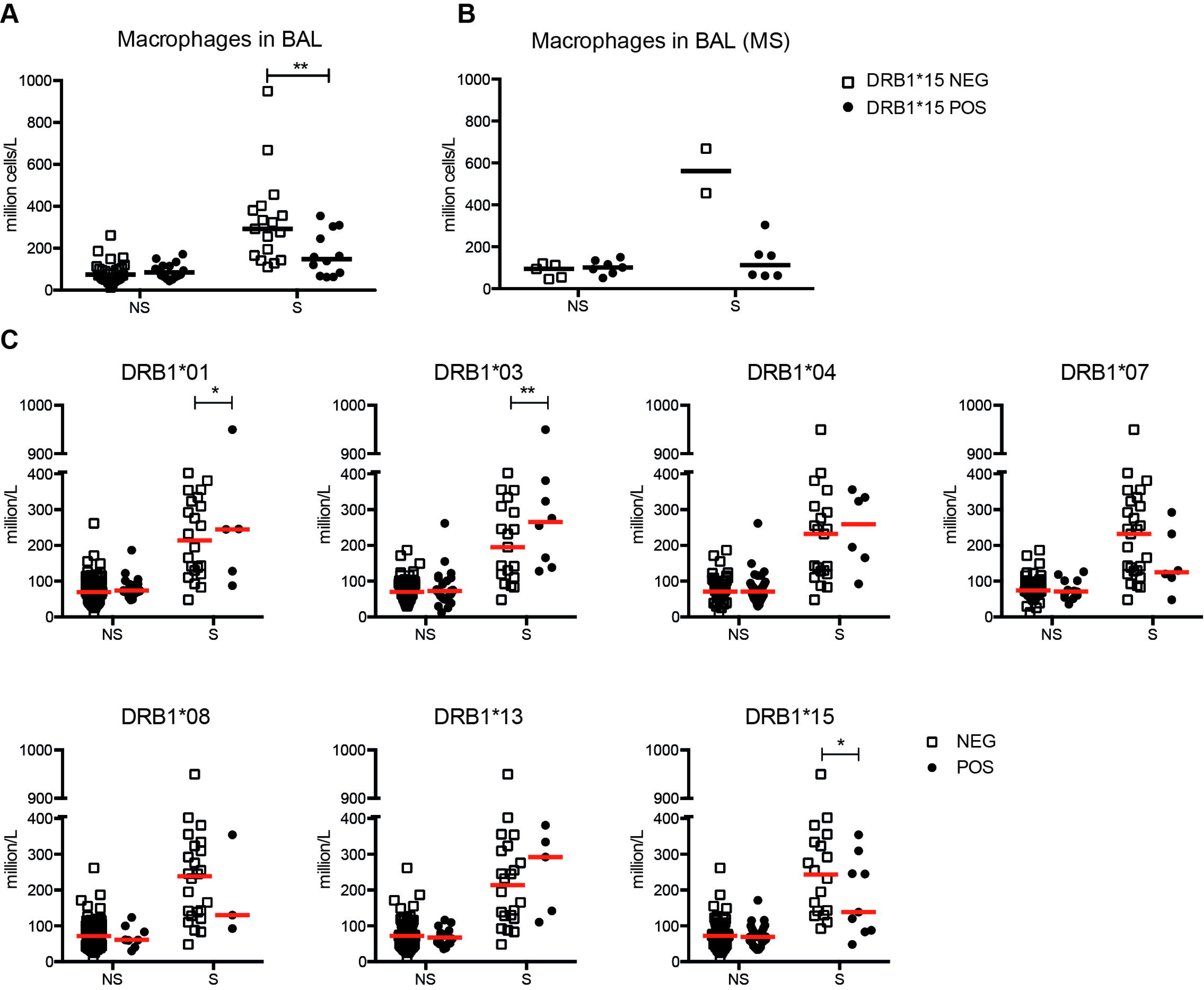
HLA-DRB1 alleles associated with macrophage concentration in the lung, in response to cigarette smoke. A) Combined analysis of healthy subjects and MS-patients, stratified on smoking status and DRB1*15 carriership. B) MS-patients only, stratified on smoking status and DRB1*15 carriership. C) Analyses of the extended cohort of healthy controls, stratified on smoking status and respective HLA-DRB1 allele. Horizontal lines indicate median, filled circles indicate carriers and open boxes indicate non-carriers, of each respective allele. Alleles with<5 carriers were not included in analysis. Statistics calculated using 2-way ANOVA, with Bonferroni post-test; ^*^: p <0.05, ^**^: p <0.01. NS: non-smokers, S: smokers

## 4 Discussion

In this first systematic investigation of the pulmonary immune system in smokers and non-smokers diagnosed with MS, we confirm that smoking considerably influences the frequency, distribution and proliferation of immune cells in the lung. The magnitude of the increase in cell concentration is associated with HLA-DRB1 alleles, in particular DRB1*15. In addition we identified an increase of pCD40L in CD4+ T-cells from the lungs of non-smokers with MS, thus further implicating the lung as an immunomodulatory organ with relevance for MS.

We identified a minor airflow obstruction in the MS-patients, compared to healthy controls, regardless of smoking status, as reflected by an overall reduction of the median FEV1/FVC ratio. For this study, reversibility of airflow limitation was not assessed, and we can therefore not further define this subclinical decline in respiratory function. This could however be an indication of respiratory muscle weakness associated with lesions in the respiratory motor pathways, as previously suggested [21, 22]. Alternatively, the observed obstruction could be related to shared mechanisms in MS and chronic obstructive pulmonary disease (COPD), as COPD patients have an increased risk for MS, independently of smoking [23].

This study confirms that smoking is a major regulator of the frequency, distribution and activation of immune cells in the lung [16, 24]. In smokers, including MS-patients, the number of alveolar macrophages is dramatically increased, and the relative distribution of alveolar macrophages and lymphocytes in BALF is shifted. Our analysis also suggests that the cellular concentration in BALF content is modified by increasing age, even though age is not associated with changes in the subsets, and other analysed characteristics, of T-cells. Little is known of the biological mechanisms underlying the increased risk for autoimmune diseases in smokers, and further analysis of cellular activation and reprogramming in the lung might identify novel pathways contributing to disease pathogenesis. Here we focused on analysis of subsets and activation state of T-cells in BALF and identified a reduction of CD4+ cells associated with smoking. Analysis of Ki-67 in T-cells in BALF indicate that both MS and smoking associate with an increase in proliferating T-cells, and we further identified an overall increase in proliferating T-cells in the lung compared to blood. We can thereby establish that T-cell proliferation is prominent in the lung, and is further increased by environmental triggers, and inflammatory processes that increase proliferation of peripheral T-cells, such as MS[25].

Interaction between CD40 on APCs, and CD40L on CD4+T-cells is essential for initiating and maintaining adaptive immune responses, including macrophage activation and licensing of dendritic cell (DCs). The interaction allows in particular DCs to functionally mature and subsequently promote activation of CD4+ and CD8+ T-cells. The expression of surface CD40L on T-cells is transient and the retention stringently regulated. The mechanisms ensuring this restricted surface expression include endocytosis mediated by CD40, and shedding of CD40L by CD40+ cells [26–28].

Two-photon microscopy studies have shown that the majority of interactions between effector/memory CD4+ T-cells and APC's *in vivo* are surprisingly brief, ranging from five to thirty minutes [29]. This time is too short to allow for *de novo* synthesis of CD40L in T-cells (detectable within 2 hours), and recent studies have indicated that intracellular stores of pCD40L in CD4+ T-cells can translocate to the cell surface within minutes and thus activate APCs, during these brief interactions [30]. As these transient encounters are suggested to dominate over longer interactions *in vivo*, the capability to rapidly mobilize CD40L from preformed stores is thus crucial for ensuring an efficient interaction between T-cells (CD40L+) and APCs (CD40+). While rapid surface mobilization of pCD40L appears to be a general characteristic of all effector and memory CD4+ T-cells, increased levels of pCD40L might be indicative of cellular activation or recent antigen encounter, as this has been demonstrated in virus-specific effector and memory CD4+ T-cells after infection [30]

Increased expression of CD40L has previously been observed in peripheral blood T-cells from MS patients, with an attenuated increase associated with IFN-β treatment [31, 32], and blocking of CD40L in EAE induced mice lead to a decrease in disease severity [33]. Here we observed a marked increase in preformed CD40L expression in pulmonary T-cells from non-smoking MS-patients, regardless of DMT status. This observation suggests that increased levels of pCD40L in MS patients is related to prior antigen exposure or changes in the lung inflammatory milieu that promotes sustained high levels of intracellular pCD40L, and could thus result in more efficient and prolonged interactions between antigen specific T-cells and APCs.

We also identified an association between HLA-genotype and the pulmonary response to smoke exposure. In both MS-patients and healthy subjects, the increase in concentration of AM associated with smoking is attenuated in HLA-DRB1*15 carriers, thus providing a first insight to a possible biological mechanism underlying the gene-environment interaction previously described [8]. In contrast, DRB1*01 and *03 is associated with a higher concentration of AM in BALF from smokers, compared to the respective non-carriers. Although the numbers of carriers are low for certain alleles, this nevertheless supports the notion that genes within, or in linkage disequilibrium with, the HLA-DRB1 region can significantly alter the immune response towards cigarette smoke. The full characterization of this altered immune response requires further studies, but could include cytokines, transcription factors, or non-coding RNAs involved in differentiation of AM precursors or recruitment of peripheral blood monocytes. The impact of the attenuated macrophage response in HLA-DRB1*15 carriers could result in a suboptimal clearance of smoke particles and a subsequently prolonged smoke-induced inflammatory response, or an altered response towards pathogens in the airways. Additional analyses of AM phenotypes and activation states will be necessary to reveal the extent of the genetic regulation of the inflammatory response induced by smoke exposure.

In summary, we here establish the lung as a target for studies of smoke-related autoimmune diseases. Further studies of smoke-induced immune responses and gene-environment interactions in the lung has the potential to define the molecular mechanisms underlying the increased risk for disease conferred by cigarette smoking and reveal novel pathways involved in MS and other autoimmune diseases.

## 5 Acknowledgements

We thank all the patients and volunteers contributing to the study. We would also like to thank Heléne Blomqvist, Margitha Dahl, and Gunnel de Forest for recruitment and management of patients and volunteers. We also thank Dr Pernilla Stridh and Dr Ingrid Kockum for assistance with HLA genotypes.

This study was supported by the Swedish Research Council, the Swedish Heart-Lung Foundation, Knut and Alice Wallenberg Foundation, Neuro Sweden, the Swedish MS-foundation, Karolinska Institutet, and Magnus Bergvall Foundation. None of the funding agencies had any role in study design, in the collection, analysis and interpretation of data, in the writing of the report or in the decision to submit the article for publication.

## References

1 H. F. McFarland, R. Martin. Multiple sclerosis: a complicated picture of autoimmunity, Nature immunology, 8 (2007) 913–919. doi:10.1038/ni1507

2 I. M. S. G. Consortium, A. H. Beecham, N. A. Patsopoulos, D. K. Xifara, M. F. Davis, A. Kemppinen, et al. Analysis of immune-related loci identifies 48 new susceptibility variants for multiple sclerosis, Nature genetics, 45 (2013). doi:10.1038/ng.2770

3 H. Westerlind, R. Ramanujam, D. Uvehag, R. Kuja-Halkola, M. Boman, M. Bottai, et al. Modest familial risks for multiple sclerosis: a registry-based study of the population of Sweden, Brain, 137 (2014) 770–778. doi:10.1093/brain/awt356

4 C. O'Gorman, S. A. Broadley. Smoking and multiple sclerosis: evidence for latitudinal and temporal variation, Journal of neurology, 261 (2014) 1677–1683. doi:10.1007/s00415-014-7397-5

5 H. Kallberg, B. Ding, L. Padyukov, C. Bengtsson, J. Ronnelid, L. Klareskog, et al. Smoking is a major preventable risk factor for rheumatoid arthritis: estimations of risks after various exposures to cigarette smoke, Annals of the Rheumatic Diseases, 70 (2011) 508–511. doi:10.1136/ard.2009.120899

6 A. K. Hedström, E. Sundqvist, M. Bäärnhielm, N. Nordin, J. Hillert, I. Kockum, et al. Smoking and two human leukocyte antigen genes interact to increase the risk for multiple sclerosis, Brain, 134 (2011) 653–717.

7 A. K. Hedström, M. Bäärnhielm, T. Olsson, L. Alfredsson. Exposure to environmental tobacco smoke is associated with increased risk for multiple sclerosis, Multiple sclerosis, 17 (2011) 788–793. doi:10.1177/1352458511399610

8 A. K. Hedström, I. L. Bomfim, L. F. Barcellos, F. Briggs, C. Schaefer, I. Kockum, et al. Interaction between passive smoking and two HLA genes with regard to multiple sclerosis risk, International journal of epidemiology, 43 (2014) 1791–1798. doi:10.1093/ije/dyu195

9 R. Ramanujam, A.-K. K. Hedström, A. Manouchehrinia, L. Alfredsson, T. Olsson, M. Bottai, et al. Effect of Smoking Cessation on Multiple Sclerosis Prognosis, JAMA neurology, 72 (2015) 1117–1123. doi:10.1001/jamaneurol.2015.1788

10 A. K. Hedström, M. Bäärnhielm, T. Olsson, L. Alfredsson. Tobacco smoking, but not Swedish snuff use, increases the risk of multiple sclerosis, Neurology, 73 (2009) 696–701. doi:10.1212/WNL.0b013e3181b59c40

11 A. K. Hedström, J. Hillert, T. Olsson, L. Alfredsson. Nicotine might have a protective effect in the etiology of multiple sclerosis, Multiple Sclerosis Journal, 19 (2013) 1009–1013. doi:10.1177/1352458512471879

12 F. Odoardi, C. Sie, K. Streyl, V. K. Ulaganathan, C. Schläger, D. Lodygin, et al. T cells become licensed in the lung to enter the central nervous system, Nature, 488 (2012) 675–679. doi:10.1038/nature11337

13 C. H. Polman, S. C. Reingold, B. Banwell, M. Clanet, J. A. Cohen, M. Filippi, et al. Diagnostic criteria for multiple sclerosis: 2010 revisions to the McDonald criteria, Annals of neurology, 69 (2011) 292–302. doi:10.1002/ana.22366

14 H. H. Olsen, J. Grunewald, G. Tornling, C. M. Skold, A. Eklund. Bronchoalveolar lavage results are independent of season, age, gender and collection site, PLoS One, 7 (2012) e43644. doi:10.1371/journal.pone.0043644

15 M. W. Gustavsen, M. K. Viken, E. G. Celius, T. Berge, I.-L. Mero, P. Berg-Hansen, et al. Oligoclonal band phenotypes in MS differ in their HLA class II association, while specific KIR ligands at HLA class I show association to MS in general, Journal of neuroimmunology, 274 (2014) 174–179. doi:10.1016/j.jneuroim.2014.06.024

16 R. Karimi, G. Tornling, J. Grunewald, A. Eklund, C. M. Skold. Cell recovery in bronchoalveolar lavage fluid in smokers is dependent on cumulative smoking history, PLoS One, 7 (2012) e34232. doi:10.1371/journal.pone.0034232

17 A. L. Peters, L. L. Stunz, G. A. Bishop. CD40 and autoimmunity: the dark side of a great activator, Seminars in immunology, 21 (2009) 293–300. doi:10.1016/j.smim.2009.05.012

18 M. Ostadkarampour, A. Eklund, D. Moller, P. Glader, O. C. Höglund, A. Lindén, et al. Higher levels of interleukin IL - 17 and antigen - specific IL - 17 responses in pulmonary sarcoidosis patients with Löfgren's syndrome, Clinical & Experimental Immunology, 178 (2014) 342–352. doi:10.1111/cei.12403

19 M. Abo Al Hayja, A. Eklund, J. Grunewald, J. Wahlstrom. Reduced expression of peroxisome proliferator-activated receptor alpha in BAL and blood T cells of non-lofgren's sarcoidosis patients, J Inflamm, 12 (2015) 28. doi:10.1186/s12950-015- 0071-6

20 K. M. Ahlgren, T. Ruckdeschel, A. Eklund, J. Wahlstrom, J. Grunewald. T cell receptor-Vbeta repertoires in lung and blood CD4+ and CD8+ T cells of pulmonary sarcoidosis patients, BMC Pulm Med, 14 (2014) 50. doi:10.1186/1471-2466-1450

21 B. Buyse, M. Demedts, J. Meekers, L. Vandegaer, F. Rochette, L. Kerkhofs. Respiratory dysfunction in multiple sclerosis: a prospective analysis of 60 patients, The European respiratory journal, 10 (1997) 139–145.

22 S. C. Smeltzer, J. H. Skurnick, R. Troiano, S. D. Cook, W. Duran, M. H. Lavietes. Respiratory function in multiple sclerosis. Utility of clinical assessment of respiratory muscle function, Chest, 101 (1992) 479–484. doi:10.1378/chest.101.2.479

23 A. Egesten, L. Brandt, T. Olsson, F. Granath, M. Inghammar, C.-G. G. Löfdahl, et al. Increased prevalence of multiple sclerosis among COPD patients and their first-degree relatives: a population-based study, Lung, 186 (2008) 173–178. doi:10.1007/s00408-008-9081-y

24 H. Forsslund, M. Mikko, R. Karimi, J. Grunewald, Å. M. Wheelock, J. Wahlström, et al. Distribution of T-Cell Subsets in BAL Fluid of Patients With Mild to Moderate COPD Depends on Current Smoking Status and Not Airway Obstruction, Chest, 145 (2014) 711–722. doi:10.1378/chest.13-0873

25 D. A. Duszczyszyn, J. D. Beck, J. Antel, A. Bar-Or, Y. Lapierre, V. Gadag, et al. Altered naive CD4 and CD8 T cell homeostasis in patients with relapsing-remitting multiple sclerosis: thymic versus peripheral (non-thymic) mechanisms, Clinical and experimental immunology, 143 (2006) 305–313. doi:10.1111/j.1365-2249.2005.02990.x

26 M. J. Yellin, K. Sippel, G. Inghirami, L. R. Covey, J. J. Lee, J. Sinning, et al. CD40 molecules induce down-modulation and endocytosis of T cell surface T cell-B cell activating molecule/CD40-L. Potential role in regulating helper effector function, J Immunol, 152 (1994) 598–608.

27 D. Graf, S. Muller, U. Korthauer, C. van Kooten, C. Weise, R. A. Kroczek. A soluble form of TRAP (CD40 ligand) is rapidly released after T cell activation, Eur J Immunol, 25 (1995) 1149–54. doi:10.1002/eji.1830250639

28 C. van Kooten, C. Gaillard, J. P. Galizzi, P. Hermann, F. Fossiez, J. Banchereau, et al. B cells regulate expression of CD40 ligand on activated T cells by lowering the mRNA level and through the release of soluble CD40, Eur J Immunol, 24 (1994) 181–92. doi:10.1002/eji.1830240402

29 C. D. Allen, T. Okada, H. L. Tang, J. G. Cyster. Imaging of germinal center selection events during affinity maturation, Science, 315 (2001) 528–531. doi:10.1126/science.1136136

30 Y. Koguchi, T. J. Thauland, M. K. Slifka, D. C. Parker. Preformed CD40 ligand exists in secretory lysosomes in effector and memory CD4+ T cells and is quickly expressed on the cell surface in an antigen-specific manner, Blood, 110 (2001) 2520–2521. doi:10.1182/blood-2001-03-081299

31 N. Teleshova, W. Bao, P. Kivisäkk, V. Ozenci, M. Mustafa, H. Link. Elevated CD40 ligand expressing blood T-cell levels in multiple sclerosis are reversed by interferon-beta treatment, Scandinavian journal of immunology, 51 (2000) 312–320. doi:

32 J. Jensen, M. Krakauer, F. Sellebjerg. Increased T cell expression of CD154 (CD40- ligand) in multiple sclerosis, European journal of neurology, 8 (2001) 321–328. doi:10.1046/j.1468-1331.2001.00232.x

33 L. M. Howard, M. C. Dal Canto, S. D. Miller. Transient anti-CD154-mediated immunotherapy of ongoing relapsing experimental autoimmune encephalomyelitis induces long-term inhibition of disease relapses, Journal of neuroimmunology, 129 (2002) 58–65.

